# Intrinsic conformational equilibria position arrestin-2 for activation

**DOI:** 10.64898/2025.12.24.696424

**Authors:** Tucker J. Shriver, Kerem Kahraman, Mingzhe Pan, Çağdaş Dağ, Marco Tonelli, Scott A. Robson, Joshua J. Ziarek

**Affiliations:** Department of Pharmacology, Northwestern University Feinberg School of Medicine, Chicago, IL, USA; Department of Cellular and Integrative Physiology, University of Texas Health San Antonio, San Antonio TX, USA; Nanofabrication and Nanocharacterization Center for Scientific and Technological Advance Research (n2STAR), Koç University, Istanbul, Turkiye; Koç University Isbank Center for Infectious Diseases (KUISCID), Koç University, Istanbul, Türkiye; National Magnetics Resonance Facility at Madison, Department of Biochemistry, University of Wisconsin-Madison, Madison, WI, USA

**Keywords:** beta-arrestin, conformational equilibria, pre-activation, interdomain rotation, dynamics, thermodynamics of activation, GPCR signaling adaptor

## Abstract

Arrestins regulate G protein–coupled receptor (GPCR) signaling by undergoing large-scale conformational rearrangements, yet the solution-state equilibria that underlie arrestin pre-activation remain poorly defined. Here, we use methyl-specific nuclear magnetic resonance spectroscopy, temperature-dependent chemical shift analysis, and relaxation measurements to characterize the intrinsic conformational landscape of full-length human arrestin-2 in solution. We identify two distinct equilibria with separable thermodynamic and kinetic signatures. A slow, enthalpically-favored process sensed by interdomain isoleucines I241 and I317 populates an active-like, interdomain-twisted conformation at physiological temperatures. In parallel, a faster, globally distributed equilibrium consistent with C-terminal tail release exhibits opposing thermodynamic behavior. Dynamic analyses reveal localized rigidification in the active-like minor states despite arrestin’s overall flexibility, while backbone relaxation data indicate widespread μs–ms conformational exchange. Together, these results demonstrate that arrestin-2 intrinsically samples activation-relevant conformations in the absence of binding partners, providing a solution-state framework for arrestin pre-activation and signaling competence.

## INTRODUCTION

Arrestins are multifunctional adaptor proteins that decode the activation state of G protein–coupled receptors (GPCRs) to regulate signaling, desensitization, and trafficking. Their ability to bind phosphorylated GPCRs, sterically block G protein coupling^1,2^, and recruit diverse effector proteins requires coordinated conformational rearrangements throughout the arrestin fold. Arrestins function as dynamic signaling hubs, recognizing more than 800 GPCRs in addition to numerous non-GPCR partners^3^, and must therefore accommodate a broad repertoire of activation states and phosphorylation patterns. Their capacity to interpret such biochemical diversity arises from an intrinsic conformational plasticity that allows them to transition between distinct structural and dynamical regimes. Structural studies have established that arrestin activation involves a series of large-scale rearrangements, including dissolution of the polar core, displacement of the C-terminal tail, remodeling of receptor-interacting loops, and a characteristic ~15–20° interdomain rotation between the N- and C-terminal β-sandwich domains^4^. The first X-ray free-electron laser (XFEL) crystal structure of a rhodopsin–arrestin complex revealed how these motions stabilize an active-like conformation capable of engaging both phosphorylated receptor tails and the receptor core^5^. Subsequent structures of GPCR complexes and phosphopeptide-bound arrestins demonstrated that activation does not produce a single discrete conformer but instead populates related active-like states with differing loop geometries, β-strand registers, and interdomain twist angles^6–8^. Together, these structures demonstrate that arrestin activation does not yield a single state but rather a spectrum of conformations modulated by receptor engagement, phosphorylation patterns, and membrane context.

Receptor phosphorylation is a dominant determinant of how arrestins move across their conformational landscape. Distinct “phosphorylation barcodes” generated by GPCR kinases guide arrestin into functionally selective conformations^9,10^. Using ^19^F NMR spectroscopy with site-specifically labeled residues, Yang et al. showed that different phosphopeptides stabilize distinct arrestin-1 conformational ensembles, each associated with selective effector engagement such as clathrin binding or Src activation^11^. More recently, Isaikina *et al.* identified a consensus pXpp phosphorylation motif that specifically drives arrestin-2 toward an active-like state resembling the vasopressin receptor peptide complex, stabilizing the finger loop, lariat loop, and interdomain interface in a coordinated manner^12^. Their work demonstrates that phosphorylation motifs remodel the energetic landscape of arrestin activation in a modular, sequence-dependent fashion. Membrane phosphoinositides provide a second layer of allosteric regulation. PI(4,5)P₂ and related lipids modulate arrestin’s transition between tail-engaged and fully engaged complexes^13^, enhance receptor–arrestin stability^14^, and reshape the conformational preferences of the C-loop and finger loop^15^. Janetzko et al. showed that PI(4,5)P₂ can be required for productive GPCR–arrestin complex formation, tuning the lifetime and topology of arrestin engagement in a receptor-dependent manner^13^. These findings highlight arrestin activation as a combinatorial process governed by receptor phosphorylation, membrane composition, and receptor conformational state.

Despite these structural and biochemical advances, significant gaps remain in our understanding of arrestin’s *solution-state* conformational equilibria. Although static structures capture basal, preactivated, and fully engaged poses, multiple loop elements, the C-tail, and interdomain cleft regions are frequently unresolved or variably ordered, indicating substantial internal flexibility. Moreover, solution-phase biophysical approaches – including DEER^16^, BRET^17^, and HDX-MS^18,19^ – reveal that arrestins access additional conformational states not apparent in crystallographic snapshots and that these states are redistributed by phosphorylation and receptor subtype. The relationships among these dynamic states, their thermodynamic underpinnings, and their mechanistic connections to active structures remain incompletely defined. Prior methyl-specific NMR studies reported that two interdomain isoleucines in arrestin-2, I241 and I317, undergo slow conformational exchange^20^, raising the possibility that these probes directly monitor the interdomain rotation that accompanies activation. However, the thermodynamics, kinetics, and local dynamical characteristics of this exchange, and how it relates to faster motions distributed across the protein, have not been delineated.

Here, we combine temperature-dependent methyl chemical shift perturbations, van’t Hoff analysis, ZZ-exchange spectroscopy, semiquantitative methyl order parameters, and categorical ^15^N relaxation to dissect the conformational equilibria of cysteine-less^21^, full-length human arrestin-2 (hArr2) in solution. We show that I241 and I317 report a shared slow, enthalpically favored equilibrium whose minor state corresponds to the interdomain-twisted active conformation observed in GPCR and phosphopeptide structures. We further identify a second, distinct fast-exchange process present across multiple isoleucines and demonstrate widespread μs–ms backbone dynamics that shape the basal arrestin ensemble. Together, these findings unify structural models of receptor- and phosphorylation-driven activation with a solution-state description of arrestin’s dynamic energy landscape and establish methyl probes as powerful intrinsic reporters of activation-linked interdomain motions.

## RESULTS AND DISCUSSION

### Interdomain Isoleucine Probes Reveal a Shared Slow Conformational Equilibrium

Shiraishi and colleagues used mutagenesis to assign two distinct resonances each to the C^δ1^ methyl groups of I241 and I317^20^. The presence of two resolved resonances for a single methyl group indicates that it exchanges (k_ex_) between two distinct environments on a timescale much slower than the frequency difference (i.e. k_ex_ << Δω)^22^. The ^13^C^δ1^ chemical shift difference between the major and minor resonances of I241 and I317 is approximately 0.3 ppm; on a 600 MHz (14.1 T) spectrometer, this corresponds to Δν ≈ 45 Hz (Δω = 2πΔν ≈ 284 s^−1^), which yields an upper bound k_ex_ ≲ 1.4 × 10^2^ s^−1^. I241 and I317 are located at the interface between arrestin’s N- and C-terminal beta-sandwich domains, which rotate 10-20° relative to each other upon GPCR complex formation (Fig. 1A).^23^ We therefore hypothesized that the two slowly exchanging resonances reflect distinct values of the interdomain twist angle corresponding to different activation states. For both residues, the chemical shift difference between the major and minor resonances lies primarily along the ^13^C dimension (Fig. 1B). Methyl ^13^C chemical shifts are well-established reporter of χ2 rotameric averaging^24^, with increased population of the trans rotamer producing downfield shifts. Consistent with this behavior, the minor states of both residues populate the *trans* χ2 rotamer approximately 5% more than the major states (Fig. 1B). To relate these local side chain signatures to global structural rearrangements, we analyzed isoleucine χ2 conformations and interdomain twist angles across all available arrestin crystal structures with sufficient electron density resolution (Fig. 1C). Whereas the I317 χ2 remains *trans* in all structures (data not shown), consistent with its limited conformational variability in solution, the I241 ^13^C^δ1^ adopts a *trans* conformation only when it’s bound to the phosphorylated C-terminal peptide of the vasopressin-2 receptor (PDB 4JQI; Fig. 1C).

**Figure 1.**
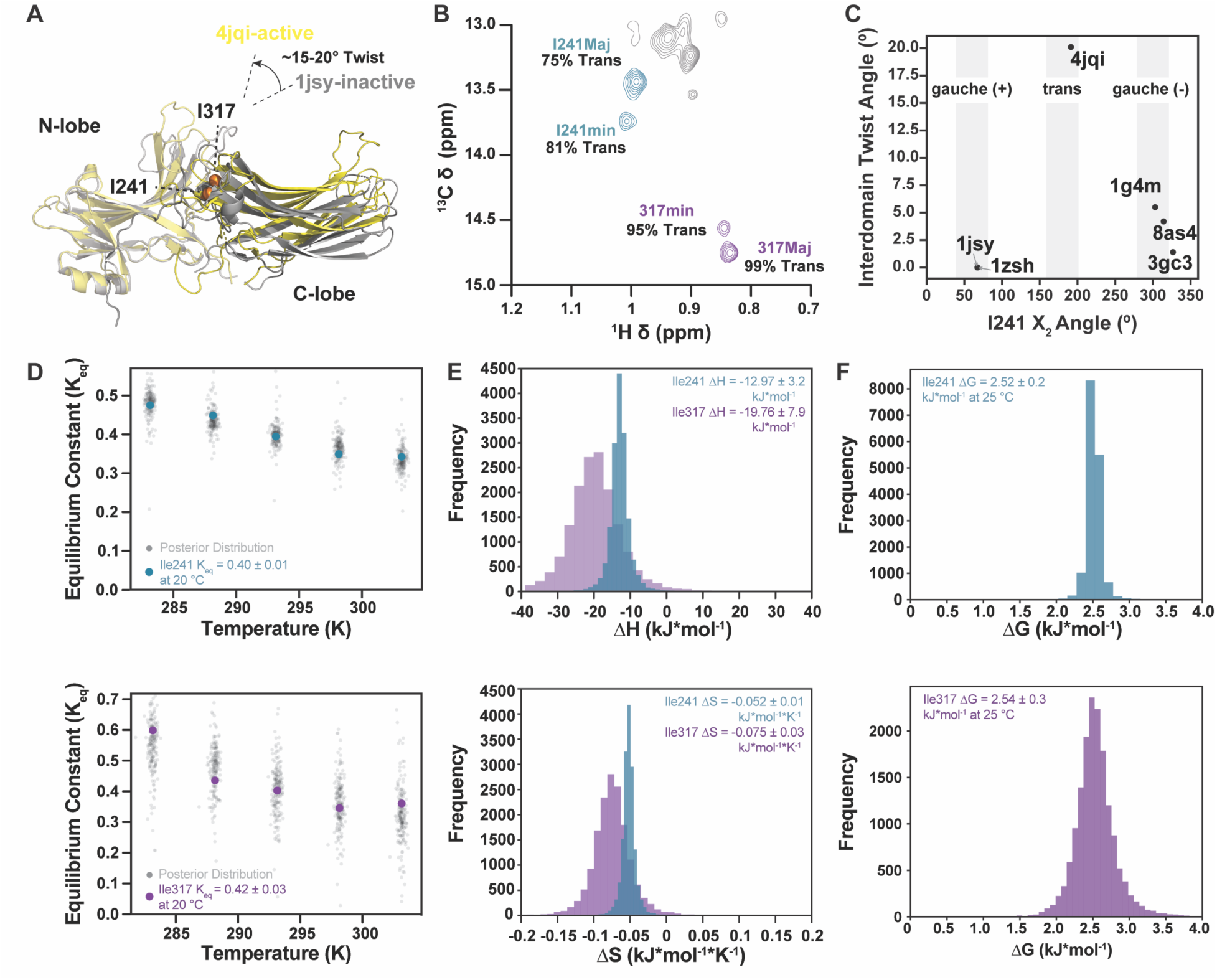
Structural and thermodynamic characterization of the slow conformational equilibrium sensed by I241 and I317. A) Overlay of basal (PDB 1JSY) and V2Rpp-bound (PDB 4JQI) Arr2 structures aligned on their N-domains, illustrating the 15-20° interdomain twist and the positions of I241 and I317 (shown as spheres) at the interdomain interface. B) Expansion of the ^1^H-^13^C HMQC spectrum showing distinct major and minor resonances for I241 and I317 C^δ1^ methyl groups. The distinct ^13^C chemical shift values reflect differing average χ2 rotameric populations, from which the percentage of trans rotamer is estimated. C) Isoleucine χ2 sidechain rotamers and interdomain twist angles taken from the Arr2 crystal structures with sufficient electron density. Structures span basal (PDB 1JSY and 8AS4), pre-activated (PDB 1G4M), in complex with hexokinase/IP6 (PDB 1ZSH), in complex with clathrin (PDB 3GC3), and active-like conformations bound to the phosphorylated C-terminal peptide from the vasopressin-2 receptor (PDB 4JQI). D) Temperature dependence of the equilibrium constant (K_eq_) between major and minor states for I241 (top) and I317 (bottom). Colored points represent experimental K_eq_ values derived from peak intensities; black curves and shaded regions denote posterior distributions from Bayesian van’t Hoff analyses. E) Posterior distributions of enthalpy (ΔH, top) and entropy (ΔS, bottom) for the conformational equilibrium obtained from the van’t Hoff analysis. F) Posterior distribution of Gibbs’ free energy (ΔG) at 298 K for I241 (top) and I317 (bottom).

Next, we acquired ^1^H-^13^C Heteronuclear Multiple Quantum Coherence (HMQC) spectra in five-degree increments from 10-30 °C, and calculated the equilibrium constant (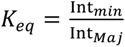) for I241 and I317 from the intensity ratio of their respective minor and major resonances (Fig. 1D). Minor state populations (P_min_) were then computed as P_min_ = Int*_min_*/(Int*_min_* + Int*_Maj_*). The temperature dependence of K_eq_ was used to estimate the change in enthalpy (ΔH) and entropy (ΔS) using the differential (i.e. non-linearized) form of the van’t Hoff equation. Both residues report an enthalpically-stabilized (ΔH < 0; exothermic) and entropically penalized (ΔS < 0) minor state (Fig. 1E). Despite difference in the magnitude of ΔH and ΔS, both residues possess similar midpoint temperatures (T_m_ = ΔH/ΔS; I241 = −23.6 °C and I317 = −9.6 °C), minor state populations (I241^min^ = 25.9% and I317^min^ = 25.7%), and Gibbs’ free energy changes (ΔG^I241^ = 2.52 ± 0.16 kJ mol^−1^; ΔG^I317^ = 2.45 ± 0.30 kJ mol^−1^) at 25 °C (Fig. 1F). Note, the negative T_m_ values represent formal extrapolations of the van’t Hoff relationship, and do not correspond to physically sampled temperatures. Taken in the context of their spatial proximity, the data suggest that both residues sense the same underlying equilibrium where the minor state is consistent with the active-like interdomain rearrangement observed in GPCR-bound structures.

### Temperature-Dependent Chemical Shift Perturbations Reveal a Distinct Fast-Exchange Equilibrium

To assess whether the other 14 isoleucine residues sense the same conformational equilibrium, we next measured the magnitude of temperature-dependent chemical shift perturbations (CSPs) within the isoleucine region from 10-30 °C (Fig. 2A). The five resonances that underwent the most substantial (≥ 0.12 ppm) combined ^1^H/^13^C CSPs (Fig. 2B) can be qualitatively binned into three categories: proton-dominated (I158), carbon-dominated (I318, I377), or mixed (I241^maj^ and I241^min^). Proton-dominated perturbations most frequently result from the ring currents of nearby aromatic groups ^25,26^ whereas ^13^C-dominated isoleucine ^13^C^δ1^ CSPs report on χ2 conformational averaging^24^. To determine the thermodynamic parameters (ΔH, ΔS, and T_m_) of each resonance, the temperature-dependent CSPs were fitted to a two-state model assuming population-weighted perturbations on the fast NMR timescale^22^ (Fig. 2C). The CSP trajectory of I158 was largely linear (data not shown), suggesting the temperature range was insufficient to sample either extreme of the equilibrium; furthermore, the extensive line-broadening of I158 is inconsistent with the assumptions of fast exchange and/or a two-state model. As the individual thermodynamic parameters of I241^maj^, I241^min^, I318, and I377 were relatively well-clustered, they were subsequently fit with global ΔH, ΔS and T_m_ values (Table 1). The consistency of individually and globally fitted parameters indicate a single, shared endothermic process (ΔH = 27.3 ± 4.8 kcal mol^−1^) that’s entropically favored (ΔS = 0.094 ± 0.02 kcal mol^−1^ K^−1^) with a T_m_ = 18.7 ± 0.6 °C. These thermodynamic signatures differ fundamentally from those obtained for the slow I241/I317 conformational exchange (Fig. 1), which exhibits ΔH < 0 and ΔS < 0 across the same temperature window. The opposite signs of enthalpy and entropy confirm that the slow and fast processes arise from distinct structural equilibria. Far-UV circular dichroism was used to estimate a global unfolding T_m_ ≈ 53.2 ± 0.2 °C, thus hArr2 undergoes a second conformational equilibrium distinct from both the I241/I317 slow exchange and unfolding (Fig. 2D).

**Figure 2.**
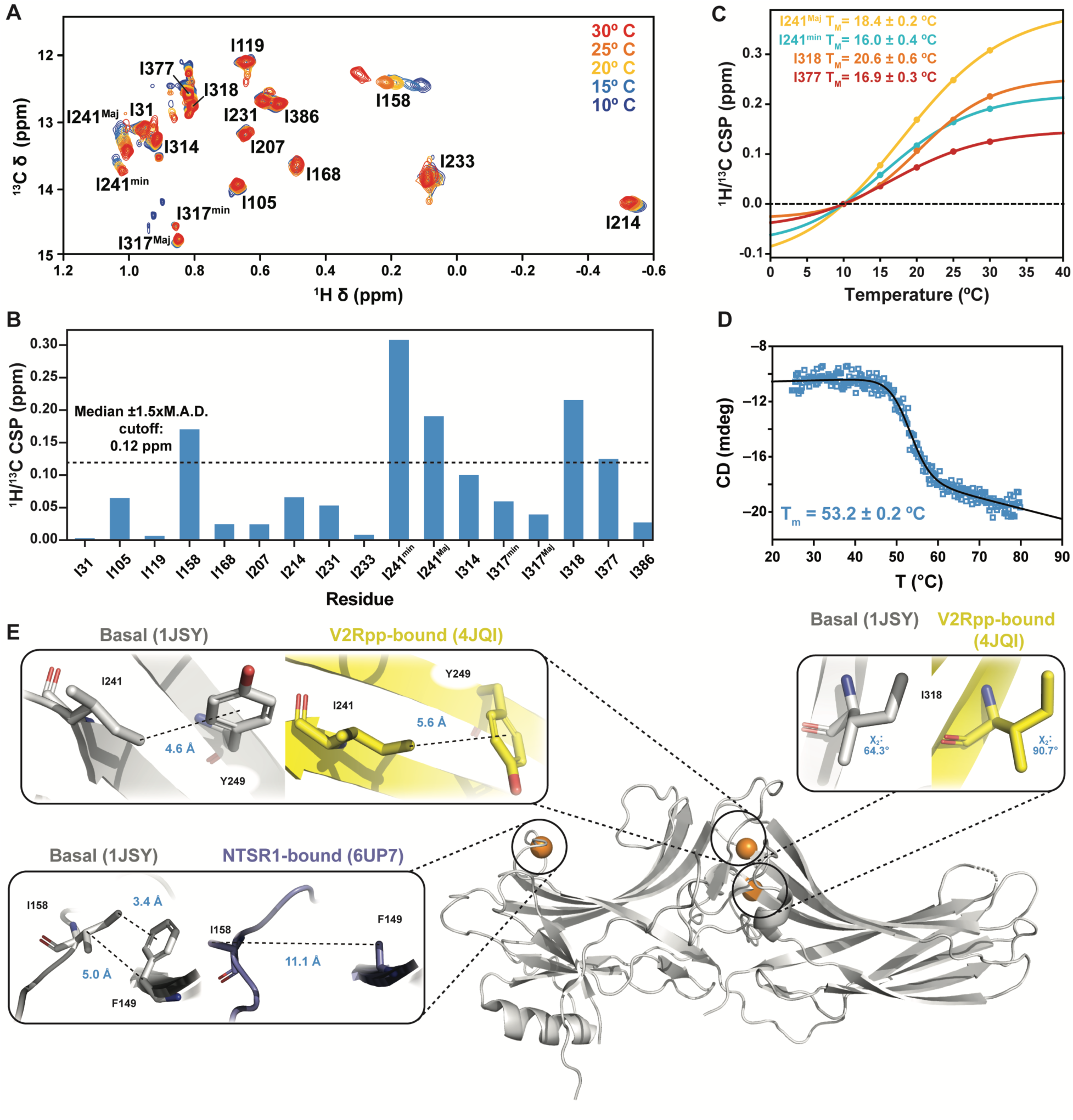
Temperature-dependent Chemical Shift Perturbation Reveal a Distinct Fast-Exchange Equilibrium in hArr2. A) Series of ^1^H-^13^C HMQC spectra collected from 10-30 °C showing temperature-dependent chemical shift perturbations (CSPs) for isoleucine C^δ1^ groups. B) Combined (^1^H/^13^C) CSPs for all isoleucines. The dotted line marks the 0.12 ppm threshold (median + 1.5x median average deviation) used to select resonances for thermodynamic analysis. C) Two-state fits of CSPs > 0.12 ppm. Fitted thermodynamic parameters are reported in Table 1. D) Far-UV circular dichroism thermal unfolding curve of hArr2, demonstrating that the CSP-derived transition (globally fit T_m_ ≈ 18.6 °C) is distinct from global unfolding (T_m_ ≈ 53.2 °C). E) Structural context for resonances exhibiting significant CSPs. Panels illustrate the local environments of I241, I318, and I158 in representative basal and active-like Arrestin-2 structures, highlighting sidechain rotamers and aromatic interactions that may underlie the observed temperature dependent shifts.

**Table 1.** Thermodynamic parameters derived from fitting temperature-dependent chemical shift perturbations. Chemical shift perturbations (CSPs) were fit individually and globally to a two-state model of the form: CSP (T) = *bottom* + (*top* − *bottom*) * (*^K^*/_1 + *K*_) where K = exp[−ΔG/(RT)] and ΔG = ΔH – TΔS. Fits yield the Gibbs’ free energy (ΔG), enthalpy (ΔH), entropy (ΔS), and midpoint temperature (T_m_ = ΔH/ΔS) for each isoleucine C^δ1^ resonance with CSPs exceeding the 0.12 ppm threshold.

| Residue | $\Delta G$ (kcal • mol<sup>-1</sup>) | $\Delta H$ (kcal • mol<sup>-1</sup>) | $\Delta S$ (kcal • mol<sup>-1</sup> • K<sup>-1</sup>) | $T_m$ (°C) |
| --- | --- | --- | --- | --- |
| I241<sup>Maj</sup> | -0.55 ± 0.03 | 24.2 ± 1.3 | 0.083 ± 0.004 | 18.4 ± 0.2 |
| I241<sup>Min</sup> | -0.86 ± 0.04 | 27.6 ± 2.0 | 0.095 ± 0.007 | 16.0 ± 0.4 |
| I318 | -0.52 ± 0.12 | 34.6 ± 5.6 | 0.118 ± 0.019 | 20.6 ± 0.6 |
| I377 | -0.74 ± 0.03 | 26.5 ± 1.6 | 0.091 ± 0.006 | 16.9 ± 0.3 |
| Global | -0.60 ± 0.10 | 27.3 ± 4.8 | 0.094 ± 0.016 | 18.6 ± 0.6 |

Again, we inspected published structures of sufficient resolution to develop mechanistic hypotheses for the temperature-dependent fast-exchange equilibria. F149 is the only aromatic residue located near I158 in any structure. In the basal state, the F149^Cβ^-I158^Cβ^ distance is 5.0 Å whereas complexation with the neurotensin receptor 1 (NTSR1) GPCR reorients I158 and increases the F149^Cβ^-I158^Cβ^ distance to 11.1 Å (Fig. 2E)^25,26^. In the basal state, the I158 C^δ1^ is positioned 3.4 Å above the plane of the aromatic ring which is consistent with the upfield (i.e. smaller) chemical shift at lower temperature. As the temperature increases, we hypothesize the GPCR-competent conformation becomes increasingly populated as reflected in the downfield (i.e. larger) chemical shift value (Fig. 2A,E). I318’s carbon-dominated CSP indicates a 10% increase in the *trans* conformation, consistent with the basal to GPCR-competent transition in published structures (~60° in PDB 1JSY vs ~90° in PDB 4JQI; Fig. 2E). I377 resides in a disordered region that has yet to be resolved in any static structure, suggesting that the *trans* conformer would dominate in solution^27^; yet surprisingly, its χ2 angle reflects a more modest *trans* rotamer (42% at 10 °C, 40% at 30 °C). Lastly, the near-equal magnitude of the I241^Maj^ and I241^min^ CSPs suggests that both states undergo a transition towards a more *trans* average rotameric conformation and a subtle reorientation relative to the nearby Y249 aromatic ring (Fig. 2E). Taken together, the thermodynamics, timescales, and residue distributions that characterize the slow and fast processes are distinct. The slow process is restricted to the interdomain region and exhibits ΔH < 0 and ΔS < 0, whereas the fast equilibrium is global and shows ΔH > 0 and ΔS > 0.

### ZZ-Exchange Establishes a Lower Bound on the Interdomain Conformational Exchange Rate

Van’t Hoff analysis of I241 and I317 major and minor resonances provided thermodynamic parameters for the conformational exchange, next we set out to determine the rate of interconversion between the two states. A first principles assessment of the ^13^C frequency difference (Δω) between major and minor resonances indicates the exchange rate (k_ex =_ k_forward_ + k_reverse_) must be slower than 63.7 s^−1^ (k_ex_ << Δω)^22^. Thus, we decided to employ the longitudinal exchange approach proposed by Tollinger and colleagues^28^ which is sensitive to time scales ranging from ~0.5 to ~50 s^−1^. Although this approach requires twice the experimental time compared to standard versions, because it requires collection of both ZZ-exchange and T1 relaxation experiments, it reduces systematic errors that could result from i) resonances that undergo additional μs-ms exchange processes, and/or ii) exchange cross peaks that are overlapped with direct correlation peaks^28^.

We recorded ^13^C ZZ-exchange and ^13^C T_1_ spectra with eight different delay times up to a maximum of 400 ms. Exchange cross peaks were not detectable even at the longest delay, which qualitatively suggests that the lifetime of each conformer is much greater than 400 ms (Fig. 3A). The major and minor peak intensities in both spectra were then fitted to four, coupled equations that yield the T_1_^Maj^ relaxation time, T_1_^min^ relaxation time, *k_Maj->min_* exchange rate constant, and *k_min->Maj_* exchange rate constant (Figure 3.B).^28^ Fits for I241 and I317 confirmed the absence of chemical exchange at the tested conditions (data not shown) and yielded T1 relaxation times that were equivalent to values determined using a simple exponential decay model. Thus, quantification of the exchange process must be limited by the T_1_ relaxation time indicating that the lifetimes greatly exceed 400 ms; these results establish only a lower bound on the timescale and do not permit quantification of the exchange rates beyond confirming the slow-exchange behavior. The median T1 relaxation time for all fifteen isoleucine resonances was 0.346 s with three (I158, I314, and I377) diverging by at least two median absolute deviations (MADs) (Fig. 3D). Mapping the T_1_ relaxation times onto the Alphafold3^29^ (AF3) model of hArr2 demonstrates I158, I314 and I377 are all located at flexible regions were relaxation times greater than the median value are expected^22^ (Fig. 3E). The T_1_ values for I241 and I317 minor states are statistically distinct from their corresponding major states, although both are within one MAD of the median, suggesting unique ps-ns dynamics in the two conformers.

**Figure 3.**
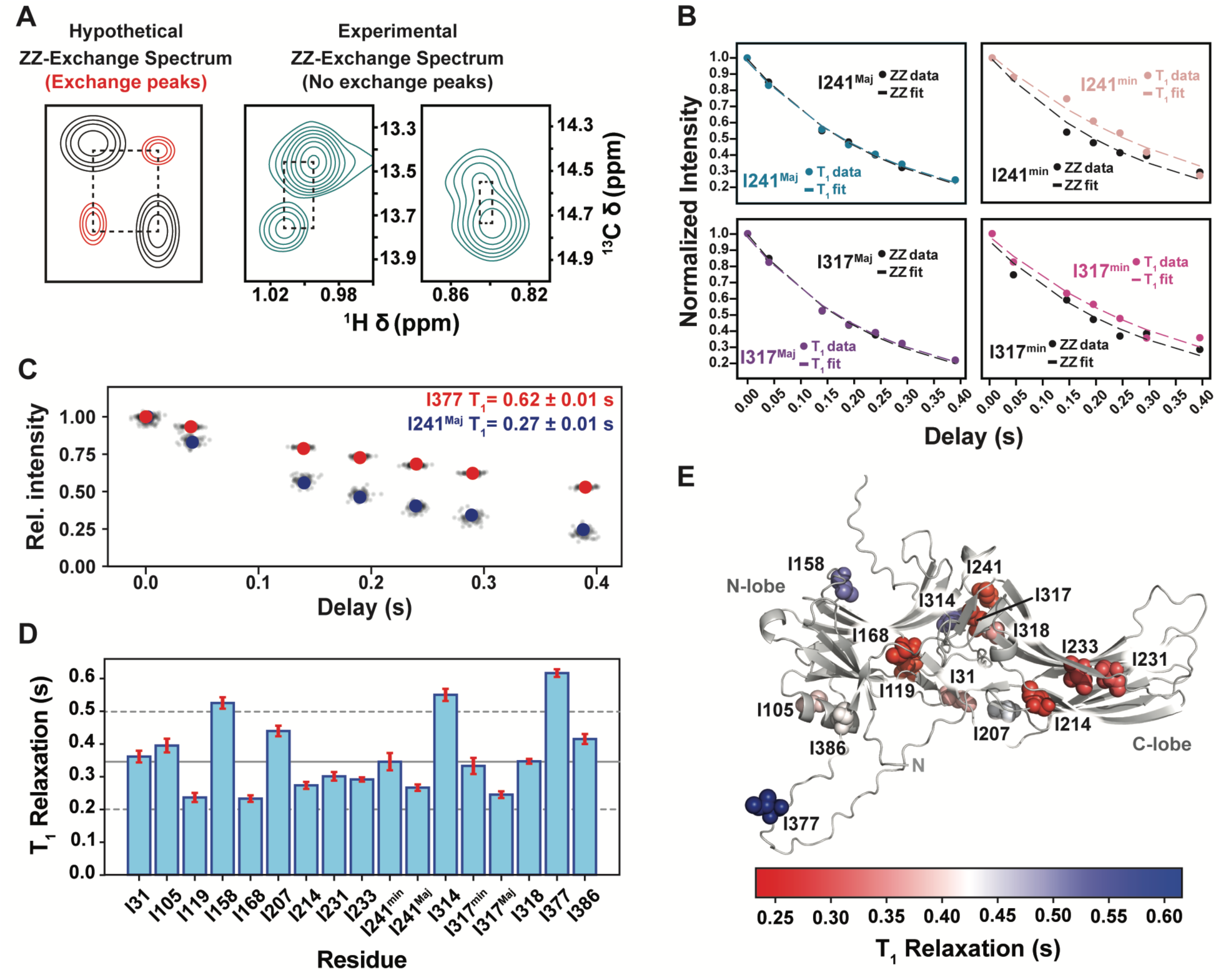
ZZ-Exchange Establish a Lower Bound on the Interdomain Conformational Exchange Rate. A) Theoretical ^13^C ZZ-Exchange spectrum (left) and experimental spectra collected at 400 ms mixing delay (right). No appreciable exchange peaks were observable at the longest delay (400 ms), indicating that the interdomain exchange process is slower than the detection limit of the experiment. B) Global fitting of the T_1_ relaxation and ZZ-exchange peak intensities for I241 and I317. Although T_1_ values are well determined, no measurable exchange contribution is detected, consistent with lifetimes substantially longer than 400 ms. C) Representative T_1_ relaxation curves for I377 (red) and I241^Maj^ (blue) with posterior-derived values shown in black. D) T_1_ relaxation times (with uncertainties) for all isoleucine residues. The median T_1_ = 0.346 s with a median absolute deviation (MAD) of 0.074 s. I158, I314, and I377 T_1_ relaxation times exceed 2 x MAD from the median. Major and minor states of I241 (p = 0.0016) and I317 (p = 0.00027) exhibit statistically distinct T_1_ values. E) T_1_ values mapped onto the AlphaFold3-derived structure of apo hArr2.

### Methyl Order Parameters Reveal Increased Rigidity in the Minor State

While the I241 and I317 ^13^C^δ1^ minor state T_1_ values lie near the median, the major states’ shorter T_1_ relaxation times (faster R_1_ relaxation rates) are suggestive of distinct mobility on the ps-ns timescale such as increased amplitude of the C^δ1^-C^γ1^ bond librations, acceleration of χ2 rotamer interconversion, slowing of local loop/backbone fluctuations, and/or increased intermolecular packing/contacts^30^. As deconvolution of these multiple mechanistic models is nontrivial, we decided to characterize isoleucine ^13^C^δ1^ motions using Lipari-Szabo “model-free” approach^31^; this formalism simplifies the description of fast internal motions to their amplitude (aka squared generalized order parameter; *O^2^*) and effective correlation time (τ_e_). In the specific case of methyl groups, the intramethyl rotameric interconversion correlation time is so fast that its role can be effectively ignored^32^. Thus, we focused our analysis on the methyl axial order parameter squared (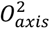), which quantifies how rigidly a methyl group’s rotation axis (C^δ1^-C^γ1^ bond) is oriented, ranging from 1 for completely rigid to 0 for fully isotropic motion. In the specific case of isoleucine 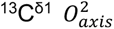, high order parameters reflect sidechains that fluctuate within a single χ2 rotameric well whereas smaller 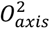 values reflect an increasing frequency of interconversions^33^.

We used the semiquantitative approach developed by Tugarinov and Kay to estimate the methyl axial order parameter^34^. Briefly, assuming HSQC and HMQC spectra were collected with identical acquisition parameters on a protein with a known overall rotational correlation time (τ_c_), this approach reasons that the intensity ratio of an HMQC and HSQC cross peak (I_HMQC_/I_HSQC_) is dependent on two variables: 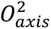 and the distance between the methyl protons and remote protons 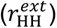. Paired HMQC/HSQC experiments and the multi-quantum transverse relaxation experiment were collected on a [U-^2^H;^13^CH_3_-ILVA]-hArr2 sample at 298.15 K. The I_HMQC_/I_HSQC_ ratios are shown in Figure 4A and fall within reasonable, previously described ranges^34^. The interproton distances were estimated from fits of the slow transverse relaxation rate of multi-quantum coherences (Fig. 4B).

**Figure 4.**
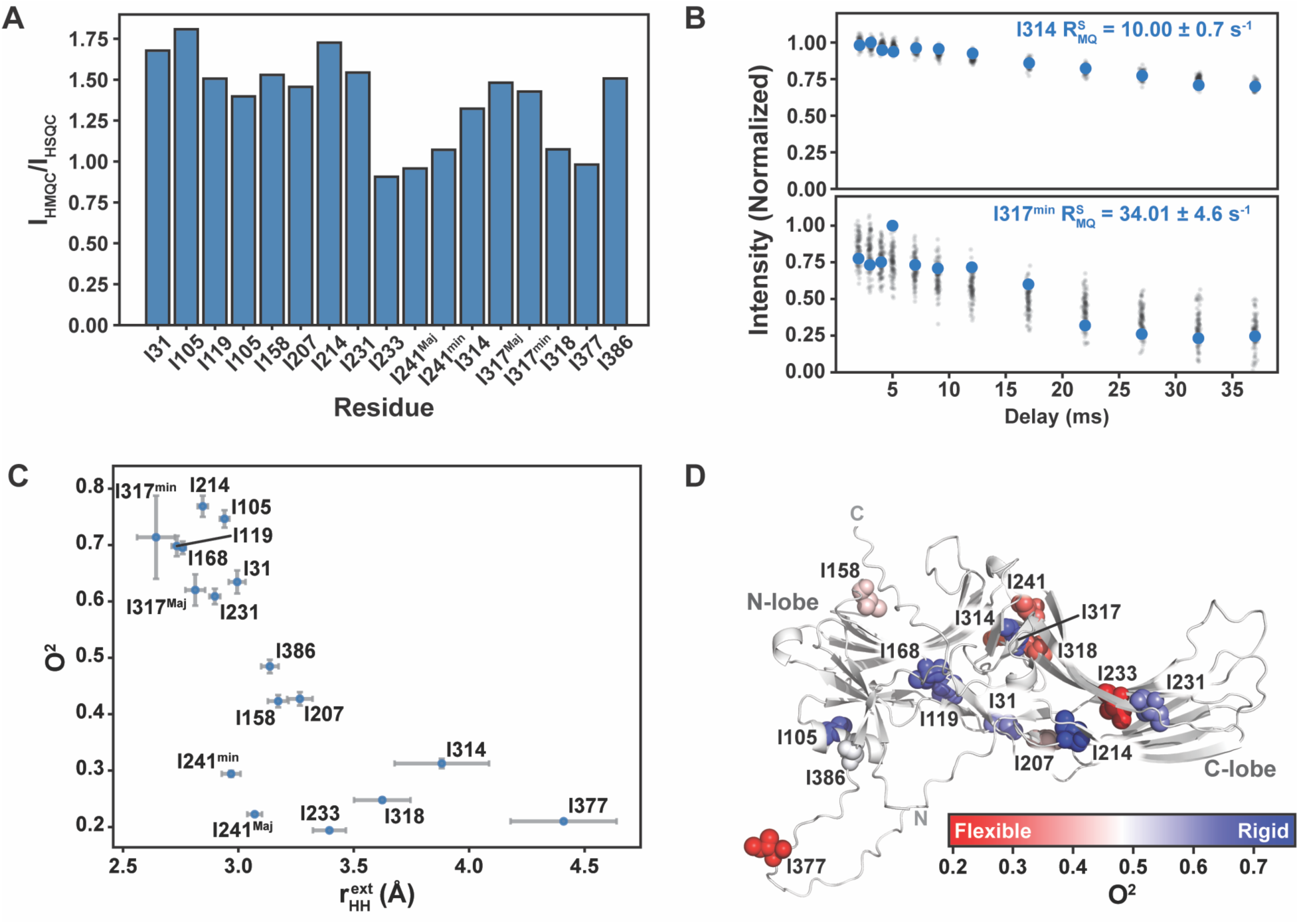
Methyl Order Parameters Reveal Increased Rigidity in the Minor States of I241 and I317. A) Ratios of ^1^H-^13^C HMQC to peak intensities (I_HMQC_/I_HSQC_) for all isoleucine methyl groups. These semiquantitative ratios provide initial estimates of methyl axial order parameters (O^2^). B) Representative multiquantum slow relaxation rates (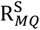) for I314 (upper) and I317^min^ (lower) shown in blue with posterior-derived points in black. C) Estimated methyl axial order parameters (O^2^) plotted as a function of the corresponding intermethyl proton distance (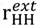). Error bars reflect posterior uncertainty in O^2^. D) Methyl axial order parameters (O^2^) mapped onto the AlphaFold3-derived apo hArr2 structure. Residues with higher O^2^ values are interpreted as more rigid (reduced χ2 rotameric interconversion), while lower values reflect increased local flexibility.

The resulting 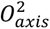 values are plotted versus their respective 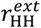 distances (Fig. 4C) and mapped onto the basal-state AF3 model (Fig. 4D). At first approximation, lower order parameters (i.e. higher flexibility; frequent rotameric interconversion) would be anticipated primarily for surface-exposed residues (such as I314 and I318) or those located in flexible regions (e.g. I377). Therefore, it’s a bit surprising that the I233 ^13^C^δ1^, which is oriented toward the domain interior, has an 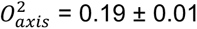 while its immediate neighbor I231 has an 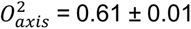 especially considering the similarity of their T_1_ values (Fig. 3D). This may reflect differences in their respective T_2_ transverse relaxation, which dominate the methyl axial order parameter^32^, and could be explored in a future study. Although both I241^Maj^ and I241^min^ exhibit more flexible order parameters than the median hArr2 isoleucine, the I241^min^ resonance undergoes less rotameric fluctuations (i.e. more rigid) than I241^Maj^; the I317^min^ resonances is similarly more rigid that its counterpart (Fig. 4C) consistent with its highly trans average conformation (Fig. 1).

### Backbone Amides Exhibit Widespread μs-ms timescale conformational exchange

To expand our understanding of the types and magnitudes of hArr2 motions beyond the limited set of isoleucine probes, we calculated the ^15^N T_1_ and T_2_ relaxation times from dynamics spectra (Fig. 5A) and determined mean values for all residues with a standard error-to-mean ratio of less than 20% (Fig. 5B). T_1_ and T_2_ relaxation times were fit for a total of 287 resonances. Following data processing, 201 resonances satisfied the criterion that the fractional uncertainty (σ/μ) for both T_1_ and T_2_ be less than 0.20. The fractional uncertainty of a fitted parameter, defined as the ratio of its posterior standard deviation (σ) to its posterior mean (μ), reflects the relative precision of the estimate. We next applied the requirement that the interscan recycling delay time is ≥ 4×T_1_ relaxation time^35^, which further reduced the dataset to 91 resonances. These resonances form the subset for categorical relaxation analysis. Although resonances failing this criterion may still exhibit interesting behavior, their T1 values cannot be reliably determined without substantially increasing the recycling delay – an adjustment that would more than triple the experimental measurement time and was therefore not feasible for this study. Because our aim in this section is to assess the extent (rather than the residue-by-residue identity) of conformational exchange in the hArr2 backbone, this filtered dataset is appropriate for a first-principles classification-based analysis.

**Figure 5.**
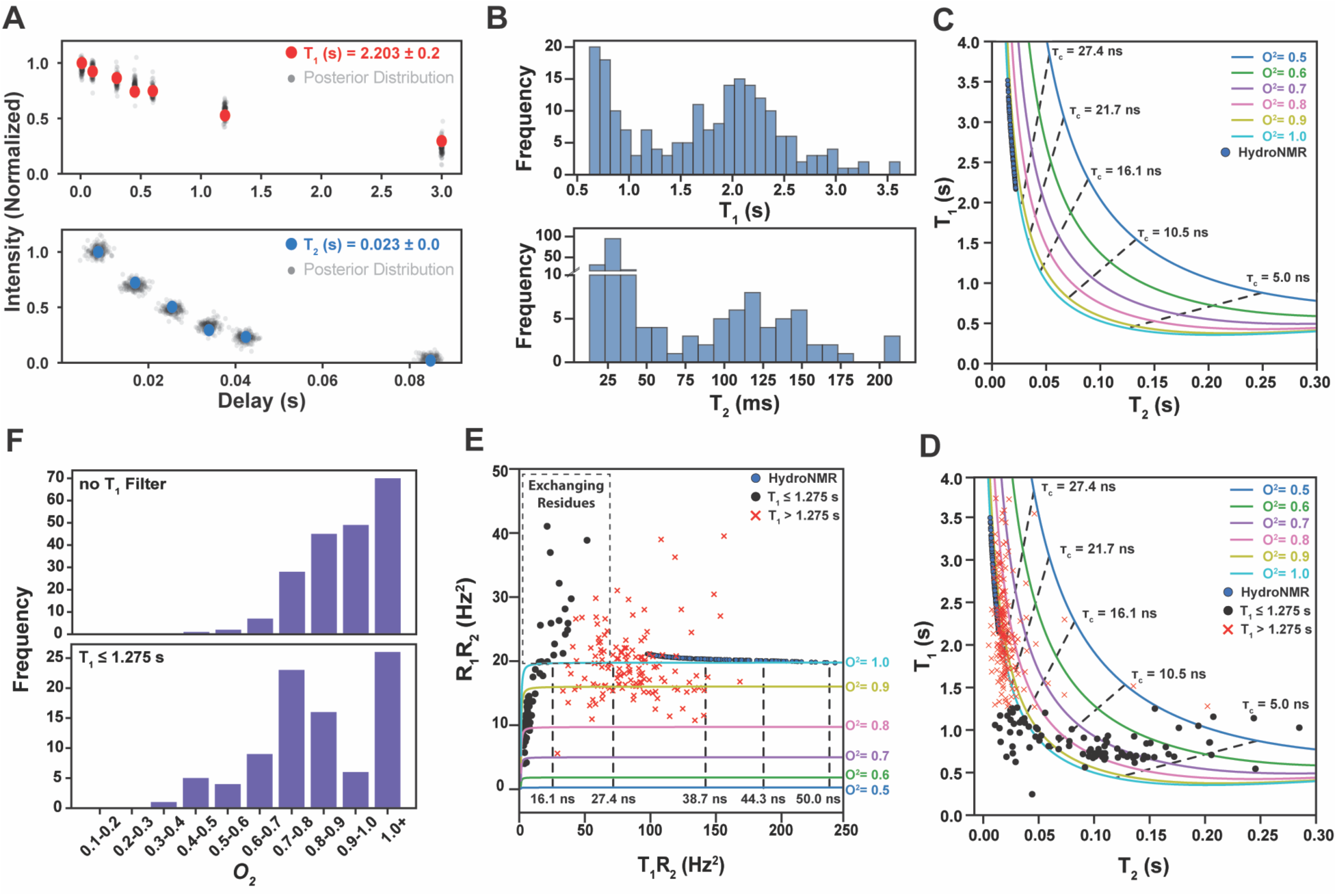
Backbone Amide ^15^N Relaxation Reveals Widespread μs-ms Conformational Exchange. A) T_1_ (upper, red) and T_2_ (lower, blue) Bayesian parameter estimations for a representative unassigned amide resonance. B) Histograms of T_1_ and T_2_ values for residues with fractional uncertainty (σ/μ) < 20%. C) Theoretical T_1_-T_2_ map showing generalized order parameter (O^2^; solid lines) and rotational correlation time (*τ_c_*; dashed lines) isolines calculated using the Lipari-Szabo model-free formalism. Blue points represent theoretical relaxation values predicted from basal-state hArr2 (PDB 8AS4) using HydroNMR assuming an axially symmetric diffusion tensor and completely rigid order parameter (O^2^ = 1). D) Experimental hArr2 ^15^N T_1_/T_2_ values overlaid on the theoretical T_1_-T_2_ map. Black points indicate resonances with sufficient recycling delay (> 4×T_1_), while red crosses indicate resonances that fail this criterion. E) Product of relaxation rates (R_1_R_2_) plotted for the same residues as in panel D. Resonances lying above the terminal O^2^ = 1 isoline indicate the presence of μs-ms conformational exchange. F) Histograms of unfiltered (upper) and filtered (lower) T_1_ values, illustrating the effect of enforcing adequate recycling delays on the distribution of relaxation parameters.

Figure 5C illustrates the theoretical ^15^N T_2_ and T_1_ isolines derived using the simplified Lipari-Szabo model-free formalism^36^. Each isoline corresponds to a particular order parameter (*O^2^*) and is cross-sectionally subdivided by overall rotational correlation times (τ_c_). Resonances that lie outside the terminal (*O^2^* = 1) isoline cannot be explained by rigid-body tumbling alone. The simplest explanations for this behavior would be that these resonances undergo additional μs-ms timescale chemical exchange or the protein possesses an anisotropic rotational diffusion tensor^37^. Plotting the product of relaxation times (R_1_R_2_) eliminates any anisotropic effects to clearly identify resonances undergoing chemical exchange^38^.

To contextualize our data, ^15^N T_1_ and T_2_ relaxation times were predicted for the basal-state hArr2 structure (PDB 8AS4) using HydroNMR^39^; the calculations employed an axially symmetric (*D_x_* ≅ *D_y_* ≠ *D_z_*) diffusion tensor and an idealized rigid generalized order parameter (*O^2^* = 1). Although theoretical, these predictions provide a benchmark for the relaxation values expected in the absence of μs-ms timescale chemical exchange (Fig. 5C). For comparison, the experimentally determined relaxation times (Fig. 5D) reveal 25 resonances lying outside the *O^2^* = 1 isoline; these same resonances also lie outside the expected region in the R_1_R_2_ transform (Fig. 5E), confirming that they undergo μs-ms conformational exchange.

Finally, the order parameter histogram derived from the interscan delay-filtered T_1_ dataset exhibits a more physically reasonable distribution than the unfiltered data (Fig. 5F). Well-ordered protein regions generally possess backbone ^15^N order parameters in the range 0.85 ≤ O^2^ ≤ 0.95^40,41^, whereas hArr2 amide residues populate the O^2^ = 0.7-0.8 bin with the highest frequency (Fig. 5F). This indicates that basal arrestin is more dynamically flexible than a canonical well-ordered globular protein, even after excluding the residues undergoing μs-ms exchange.

## CONCLUSION

This work provides a solution-state framework that directly links arrestin-2 conformational equilibria to its signaling function as a multifunctional adaptor protein^42^. Arrestins must decode diverse receptor phosphorylation patterns^9,10,12^, membrane environments^13^, and effector^21^ demands to direct signaling outcomes downstream of GPCR activation, yet how this information is encoded dynamically within the arrestin fold has remained unclear. Our results demonstrate that apo arrestin-2 constitutively samples an active-like, interdomain-twisted conformation^20,23^ that is energetically accessible under physiological conditions, positioning arrestin not as a passive responder to receptor engagement but as a preorganized signaling scaffold poised for rapid activation.

By resolving two distinct conformational equilibria with separable thermodynamic and kinetic signatures, we clarify how arrestin activation can proceed through modular, rather than strictly concerted, structural transitions. The slow exchange process reported by I241 and I317 corresponds to interdomain rotation, the hallmark rearrangement that enables arrestin to engage phosphorylated receptor tails and receptor cores in signaling-competent complexes^23^. In contrast, the fast, globally distributed equilibrium is consistent with C-terminal tail release, a prerequisite for both receptor binding^5^ and effector recruitment, including clathrin^43^, AP2^44^, and Src-family kinases^45^. The opposing thermodynamic signatures of these processes, enthalpically-favored interdomain twisting versus entropically-driven tail disengagement, support a hierarchical activation model in which distinct signaling-relevant elements of arrestin are regulated semi-independently. Importantly, our data reveal that arrestin activation is accompanied by localized rigidification rather than global stabilization. The active-like minor states sensed by I241 and I317 exhibit increased fast-timescale order despite arrestin’s overall dynamic character. This behavior is consistent with models in which arrestin leverages intrinsic flexibility to recognize diverse receptors and phosphorylation barcodes, followed by structural consolidation to transmit signals with specificity and fidelity^46,47^. The widespread μs–ms backbone dynamics observed across arrestin-2 further suggest that signaling bias and pathway selectivity may emerge from redistribution of pre-existing conformational ensembles rather than from formation of entirely new structures.

From a signaling perspective, the identification of I241 as a robust, non-perturbative sensor of interdomain activation has particular significance. This residue reports directly on a conformational transition that is central to arrestin-mediated desensitization, trafficking, and G protein–independent signaling. As such, it provides a powerful intrinsic probe for dissecting how different GPCRs, phosphorylation motifs, and membrane lipids reshape arrestin’s signaling landscape. Extending this approach to receptor-bound and membrane-associated states will be critical for determining how closely the apo active-like ensemble resembles signaling-competent complexes and how arrestin integrates multiple biochemical inputs to direct pathway-specific outcomes.

Overall, this study advances a physically grounded view of arrestin signaling in which functional specificity arises from thermodynamically and dynamically distinct conformational equilibria embedded within the basal protein ensemble. By linking interdomain rotation, tail release, and fast-timescale rigidity to signaling-relevant structural transitions, our findings provide a mechanistic bridge between arrestin dynamics in solution and its central role in GPCR signal transduction.

## MATERIALS AND METHODS

### Arrestin-2 Expression and Purification

*E. coli* (BL21-DE3) were transfected with a pET15b plasmid from the Manglik lab (UCSF). The plasmid encodes a hexahistidine-tag, 3C cleavage site followed by the cysteine-less version of full-length human Arrestin-2^21^; the primary sequence of the final protein product is below. Cells were streaked on antibiotic resistant plates and initial H_2_O-LB cultures of 5 mL were inoculated. Cells were pelleted after ~8 hours of growth and used to inoculate a 5 mL of D_2_O-LB culture overnight, which was subsequently spun down and used to inoculate a 5 mL D_2_O-2xM9 culture. These were then used to inoculate a 50 mL starter culture and eventually six 1 L D_2_O-2xM9 cultures. These 1L cultures were supplemented with ^15^NH_4_Cl and [*U*-^2^H] glucose prior to inoculation, [^1^H_3_,^13^C]-labeled methyl precursors were added (300 mg/L hydrolyzed acetolactate, 300 mg/L alanine) 1 hour prior to induction with 400 μM IPTG, and 50 mg/L ketobutyrate was added twenty-five minutes before induction. Cells were grown for 18 hours and spun down at 5000 x g. Pellets were immediately resuspended in 40 mL/L expression solubilization buffer (20 mM HEPES, 500 mM NaCl, 2 mM MgCl_2_, 10% glycerol, pH 7.5) supplemented with EDTA-free protease inhibitor (1 tablet/12 L), and 100 mg powdered DNAse and Lysozyme with stirring at 4 °C for thirty minutes. Cells were then sonicated twice for a total of 5 minutes (5 s on, 15 s off) at 35% duty cycle followed by a 5 min waiting period. Cell debris was then pelleted at 24,424 x g and the lysate run through a 0.45 μm filter before augmentation with 10 mM imidazole and loaded onto a 5 mL Cytiva HisTrapHP column at 2 mL/min. The column was washed with loading buffer (20 mM HEPES, 150 mM NaCl, 10% glycerol, pH 7.5) augmented with 8 mM ATP for 25 mL, left to rest for 15 minutes, and then washed with another 25 mL of ATP-loading buffer. The column was then washed with loading buffer until the UV absorbance was under 50 mAU and then washed with 8% elution buffer (loading buffer with 500 mM imidazole) until the absorbance was again under 50 mAU. The protein was then eluted off the column with 40% elution buffer. The protein was cleaved using 3C protease produced in our lab at 4 °C overnight with rotation. The cleavage reaction was pelleted and the supernatant run through a 0.45 um filter before diluting 1:5 with ion-exchange (IEX) A buffer (20 mM HEPES, pH 7.5) and loaded onto a 5 mL Cytiva HiTrap Q HP column again at 2 mL/min. The column was washed with 10% IEX B buffer (IEX A buffer with 500 mM NaCl) and then the protein was eluted with a 10-100% gradient over 25 minutes. Appropriate fractions (near 18 mS/cm conductivity) were concentrated using a 30 kDa spin filter, any particulate filtered out through a 0.22 um spin filter, and injected onto a Cytiva S75 prep-grade size exclusion column (SEC) using a 5 mL loop. The loop was rinsed with 20 mL (4x loop volume) of SEC buffer (20 mM sodium phosphate, 150 mM NaCl, pH 6.8) to ensure complete sample loading and 2 mL fractions were collected and then appropriately concentrated to a working concentration. The primary sequence of the final hArr2 used herein is: GPSGDKGTRVFKKASPNGKLTVYLGKRDFVDHIDLVDPVDGVVLVDPEYLKERRVYVTLTVAFRYGREDLDVLGLTFRKDLFVANVQSFPPAPEDKKPLTRLQERLIKKLGEHAYPFTFEIPPNLPSSVTLQPGPEDTGKALGVDYEVKAFVAENLEEKIHKRNSVRLVIRKVQYAPERPGPQPTAETTRQFLMSDKPLHLEASLDKEIYYHGEPISVNVHVTNNTNKTVKKIKISVRQYADIVLFNTAQYKVPVAMEEADDTVAPSSTFSKVYTLTPFLANNREKRGLALDGKLKHEDTNLASSTLLREGANREILGIIVSYKVKVKLVVSRGGLLGDLASSDVAVELPFTLMHPKPKEEPPHREVPENETPVDTNLIELDTNDDDIVFEDFARQRLKGMKDDKEEEEDGT GSPQLNNR

### Temperature-dependent Heteronuclear Multi-Quantum Coherence (HMQC) Spectral Collection and Analysis

HMQC spectra were collected using the triple resonance cryoprobe-equipped Varian 800 at MetaCyt Biomolecular NMR Lab (Indiana University, Bloomington) on a 160 μM hArr2 sample in 20 mM sodium phosphate, 150 mM NaCl pH 6.8 at 283.15, 288.15, 293.15, 298.15, 303.15 K. Spectra were processed with NMRPipe^48^ hosted on NMRbox’s remote access clusters and data was visualized in CcpNMR^49^. Spectra were batch processed with zero filling and squared sine-bell window functions for apodization. Chemical shift values were recorded for all isoleucines as a function of temperature and CSPs calculated using a scaled Euclidian distance to account for the differential ^13^C and ^1^H chemical shift ranges as follows: 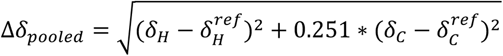, where the reference shift is the value at the lowest temperature and *δ_H_* and *δ_C_* are the proton and carbon chemical shifts respectively. These temperature-dependent CSPs were then fit to the following equation: 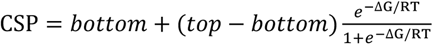 and ΔG = ΔH − TΔS.

### Circular Dichroism Melting

Sample consisted of 10 μM Arrestin 2 in 6 mM sodium phosphate, 20 mM NaCl pH 7.4. Sample was degassed for twenty minutes immediately prior to measurement. All measurements were carried out on a Jasco J-1715 circular dichroism spectropolarimeter with a Peltier temperature controller under external nitrogen flow using a 1cm pathlength quartz cuvette. All spectra were collected over a range of 190-260 nm with a scan time of 50 nm/min at a data integration rate of 1 s. Each CD trace represents the average of twenty scans. Buffer background spectra were subtracted from experimental spectra before analysis. CD values at 222 nm were monitored to determine the melting curve.

### van’t Hoff Analysis

The methyl HMQC spectra used for temperature-dependent CSP determination were used for van’t Hoff analysis. Peak heights of the major and minor peaks of I241 and I317 were recorded, and the equilibrium constant defined as 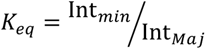. These data were then fit to the van’t Hoff equation 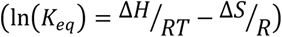 using Bayesian parameter estimation. Histograms of enthalpy and entropy values were then calculated using the posterior distributions calculated during fitting. Histograms of ΔG were then calculated at 298.15 K using these same posteriors.

### ZZ-Exchange Spectral Collection and Analysis

All spectra were collected on the same sample used for the temperature-dependent HMQCs, recorded again on the Varian 800 MHz instrument at MetaCyt, Indiana University. Interleaved T_1_ and ZZ-Exchange experiments were collected across a range of delay times up to 400 ms using the pulse programs laid out in Kloiber *et al*^50^. Relaxation decays for the major and minor peaks of I241 and I317 were fit simultaneously as a set of four, with two T_1_ decays and two ZZ decays each for the major and minor. After no appreciable K_ex_ could be determined using physically relevant definitions (*k_AB_*, *k_BA_* > 0) with non-negative forward and reverse rates, T_1_ rates were then determined for all isoleucines. Decays were fit with BPE utilizing a simple exponential model with 4000 samples and 4000 draws.

### Semiquantitative Order Parameter Spectral Collection and Analysis

All samples were collected on a 900 MHz spectrometer at NMRFAM (University of Wisconsin-Madison) using a 180 μM Arrestin 2 sample. Heteronuclear Single Quantum Coherence (HSQC) and HMQC spectra were acquired, followed by slow, multiquantum carbon relaxation spectra^51^ were collected with 12 delays up to 37 ms.

The slow multiquantum relaxation experiments were first fit using BPE to determine 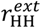 distributions. These were then used with HMQC/HSQC ratios to fit *O*^2^ values for each isoleucine residue in Arrestin 2, again using BPE with 4000 samples and 4000 draws.

### ^15^N T_1_ and T_2_ Relaxation Spectral Collection and Analysis

All data were collected on a 95 μM Arrestin 2 sample, in buffer as previous described, on the 900 MHz spectrometer at NMRFAM (University of Wisconsin-Madison). Standard amide relaxation pulse sequences were used to collect three sets of T_1_ experiments with delays of 10, 100, 300, 450, 600, 1200, and 3000 ms with carbon decoupling. T_2_ experiments were acquired with delays of 8.48, 16.96, 25.44, 33.92, 42.40, and 84.80 ms utilizing a shortened 8 ms echo. Spectra were processed with identical nmrPipe scripts containing zero filling and apodization with squared-sine bell functions.

The ^15^N-TROSY was peak picked using CCPnmr’s in-suite picking algorithm, yielding a total 333 peaks. After copying this template peak list to all T_1_ and T_2_ experiments, each peaks position was manually fit and assessed for fidelity using 1D traces, yielding 287 peaks. This set of data was then fit using a Bayesian parameter estimation model utilizing pyMC with four rounds of 4000 samples and 4000 draws to properly sample the parameter space. Signals were modeled as normalized, single exponential decays with normally distributed normalization factors, uniformly distributed relaxation constants and exponentially distributed noise used in the likelihood estimation.

The data was then filtered for unreliable fits where σ/μ ≥ 0.20 for either T_1_ or T_2_, yielding a total number of 203 usable resonances for further analysis. Paired correlation plots of T_2_ vs T_1_ and R_2_/R_1_ vs R_1_R_2_ were then overlaid with iso-order parametric curves reflecting idealized, isotropic predictions of the plots given a fixed order parameter value. These are then overlaid again with dashed lines representing iso-*τ_C_* values. Data points exceeding the O^2^ = 1 curve likely experience chemical exchange, however due to sample heating during longer delay periods, T_1_ values are inherently underestimated overestimating the number of exchanging residues.

## Author Contributions

T.J.S., S.A.R., and J.J.Z. designed research; T.J.S., K.K., C.D., M.P., M.T. performed research; T.J.S., S.A.R., and J.J.Z. analyzed data; and T.J.S., K.K., C.D., S.A.R., J.J.Z. wrote the paper.

## ACKNOWLEDGMENTS

The cysteine-less human arrestin-2 plasmid was kindly provided by Professor Aashish Manglik (University of California – San Francisco). This study made use of the National Magnetic Resonance Facility at Madison, an NIH National and Regional Resource NIH R24GM141526. Helium recovery equipment, computers, and infrastructure for data archive were funded by the University of Wisconsin-Madison, NIH R24GM141526, and National Science Foundation NSF 1946970 (NSF Mid-Scale Research Infrastructure Big Idea). The 800 MHz NMR spectrometer at Indiana University used in this research was generously supported by a grant from the Lilly Endowment.

